# Manipulation of a Cation-π Sandwich Reveals Conformational Flexibility in Phenylalanine Hydroxylase

**DOI:** 10.1101/2020.02.28.969873

**Authors:** Emilia C. Arturo, George W. Merkel, Michael R. Hansen, Sophia Lisowski, Deeanne Almeida, Kushol Gupta, Eileen K. Jaffe

## Abstract

Phenylalanine hydroxylase (PAH) is an allosteric enzyme responsible for maintaining phenylalanine (Phe) below neurotoxic levels; its failure results in phenylketonuria. Wild type (WT) PAH equilibrates among resting-state (RS-PAH) and activated (A-PAH) conformations, whose equilibrium position depends upon allosteric Phe binding to the A-PAH conformation. The RS-PAH conformation of WT rat PAH (rPAH) contains a cation-π sandwich between Phe80, Arg123, and Arg420, which cannot exist in the A-PAH conformation. Phe80 variants F80A, F80D, F80L, and F80R were prepared; their conformational equilibrium was evaluated using native PAGE, size exclusion chromatography, ion exchange behavior, intrinsic protein fluorescence, enzyme kinetics, and limited proteolysis, each as a function of [Phe]. Like WT rPAH, F80A and F80D show allosteric activation by Phe while F80L and F80R are constitutively active. Maximal activity of all variants suggests relief of a rate-determining conformational change involving Phe80. Limited proteolysis of WT rPAH in the absence of Phe reveals facile cleavage within a C-terminal 4-helix bundle that is buried in the RS-PAH tetramer interface, reflecting dynamic dissociation of the RS-PAH conformation. This cleavage is not seen for the Phe80 variants, which all show proteolytic hypersensitivity in a linker that repositions during the RS-PAH to A-PAH conformational interchange. Hypersensitivity is corrected by addition of Phe such that all Phe80 variants become like WT rPAH and achieve the A-PAH conformation. Thus, manipulation of Phe80 perturbs the conformational space sampled by PAH, increasing the propensity to sample intermediates in the RS-PAH and A-PAH interchange, which are presumed on-pathway because they can readily achieve the A-PAH conformation by addition of Phe.

## 1. INTRODUCTION

The role of phenylalanine hydroxylase (PAH, E.C. 1.14.16.1) in human biology is to maintain phenylalanine (Phe) levels below the threshold for neurotoxicity and sufficient for normal metabolism (e.g. protein, pigment and neurotransmitter biosynthesis). PAH is a ∼453 amino acid, multi-domain, multimeric protein that catalyzes the conversion of Phe to tyrosine using tetrahydrobiopterin (BH_4_), O_2_, and a non-heme iron (1). PAH dysfunction results in the most common inborn error of amino acid metabolism, phenylketonuria (PKU) (∼1:10,000 births worldwide). Without treatment, PKU has severe neurologic consequences (2). To ensure Phe homeostasis, PAH populates alternate conformations, which include a tetrameric resting state PAH (RS-PAH) conformation and a tetrameric activated PAH (A-PAH) conformation; the latter can be stabilized by allosteric Phe binding (3). Although their role in PAH allostery remains speculative (e.g. (4)), dimeric conformations are also part of the equilibrium of PAH assemblies. Some PKU-associated PAH variants have been proposed to alter the equilibrium among oligomeric forms, in particular to dysregulate the interconversion between RS-PAH and A-PAH conformations (3).

Figure 1A shows PAH segments defined by known PAH structures as well as the RS-PAH (Fig 1B) and A-PAH (Fig 1C) conformational interconversion. The sequences of rat and human PAH are compared in Fig S1. Prior to 2016, many PAH crystal structures (previously summarized in (5)) were of truncated human PAH constructs containing only segments 4 – 6, with one or more active site ligands. In the absence of segments 1-3, crystal structures show an active site lid (segment 5) in various conformations depending on the identity of active site ligands (5). Combination of structures from less severely truncated PAH constructs allowed formation of a composite homology model of the RS-PAH conformation (6), which was largely confirmed by the crystal structure illustrated in Fig 1B (5). These earlier structures did not inform on what we currently understand about the structural basis for PAH allostery, which was first proposed in 2013 (4), and established in 2016 to contain the proposed allosteric Phe binding site (7) noted by the oval in Fig 1C. PAH allostery was first described by S. Kaufman, who noted a dramatic difference in the reaction rate depending on the order of addition of reaction components (8,9). This was further correlated with a major conformational change based on the protein’s affinity for phenyl-Sepharose as a function of [Phe] (10).

**Figure 1.**
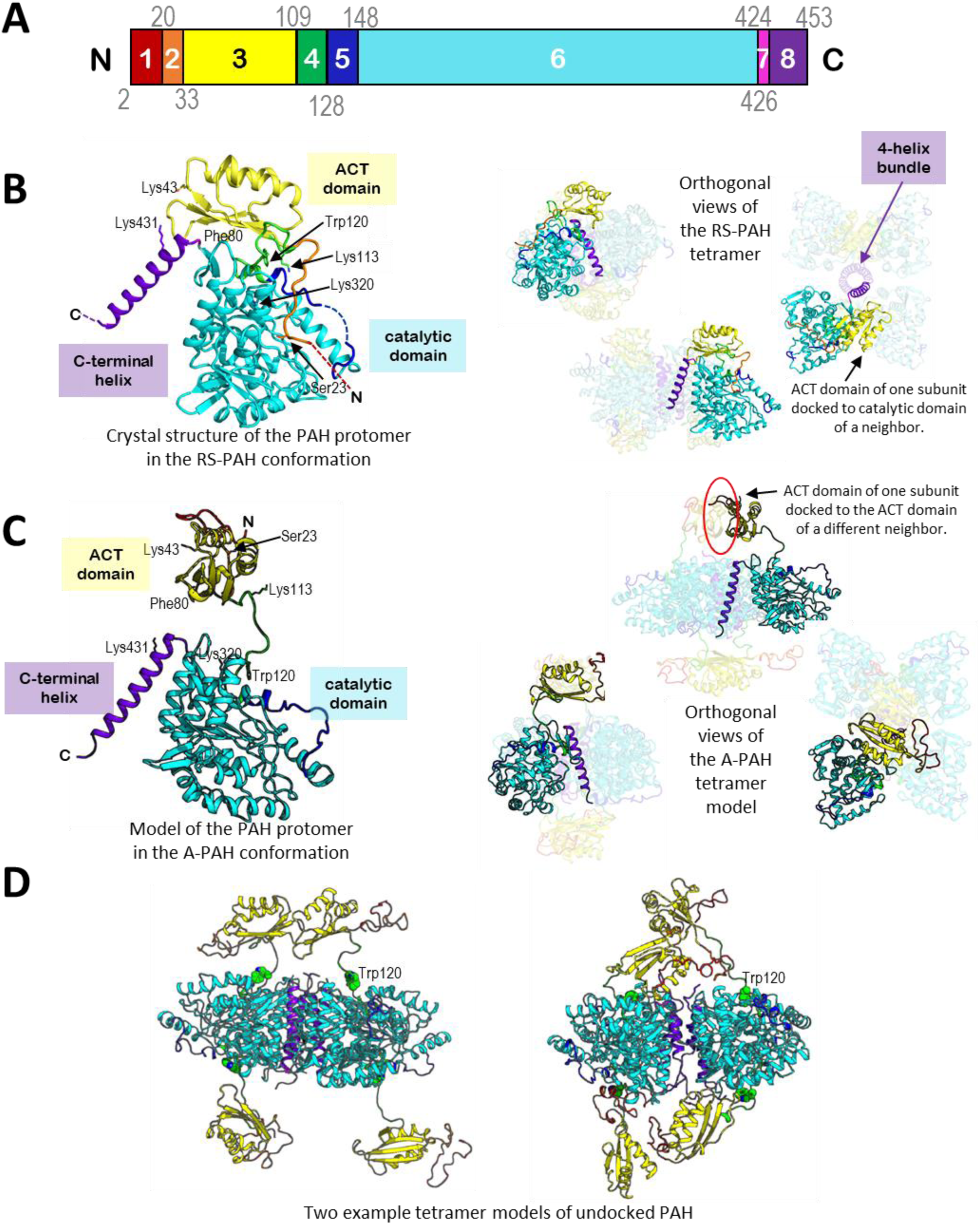
The structures of phenylalanine hydroxylase (PAH). **(A)** The domain structure of mammalian PAH is described by eight colored segments, using the same colors in the figures below. Notably, segment 1 is disordered in RS-PAH crystal structures; segment 2 contains the RS-PAH-specific autoinhibitory interaction; together these define the autoinhibitory region. With the ACT domain (segment 3), these define the regulatory domain. Segment 4 is a linker region, some or all of which repositions during the RS-PAH ⇔ A-PAH conformational interchange. Segments 5 & 6 comprise the catalytic domain; segment 5 is the active site lid. Segment 7 is a hinge whose conformation differs among the four chains in all full length RS-PAH tetramer structures (5,11,13). Segment 8 is a C-terminal helix, seen as both bent and straight, which forms a 4-helix bundle; it does not contain the classic pattern of leucine residues found in other aromatic amino acid hydroxylases (22). **(B)** An X-ray crystal structure of WT rPAH in the tetrameric RS-PAH conformation (PDB: 5DEN (5)) is shown. One subunit is enlarged on the left. Three orthogonal illustrations of the tetramer are shown on the right with three subunits faded. In this conformation, the ACT domain of each subunit is docked to the catalytic domain of a neighboring subunit along the short edge of the tetramer as illustrated far right. On the left image, we show in sticks the side chains of residues Phe80, Trp120 (important in PAH intrinsic fluorescence), and lysine cleavage sites identified by native trypsinolysis. **(C)** A recent A-PAH model is shown in the “trans” conformation (13); one subunit is enlarged on the left with the same residues depicted as part B. Three orthogonal illustration of the tetramer are shown on the right with three subunits faded. In this conformation, the ACT domain of one subunit is docked to the ACT domain of the subunit situated diagonally across the tetramer. An A-PAH-specific ACT:ACT interface is circled in red. This is where the allosteric Phe binds (7). Although the RS-PAH and A-PAH conformations differ in the position of the regulatory domain relative to the rest of the protein, both conformations contain ACT domains that are docked. All proposed A-PAH conformational models have uncertainty in the position of segments 1 + 2. **(D)** Two example models of “undocked” tetrameric PAH conformations are illustrated. The model on the left was prepared using the program SASSIE (23) and is part of a detailed SEC-SAXS analysis of the Phe80 variants (Gupta et al, in preparation). The model on the right is the result of a molecular dynamics investigation of tetrameric PAH starting from a crystal structure in the RS-PAH conformation (Ge, et al, in preparation), expanding methods we previously applied to a Phe-bound ACT domain dimer (15).

There are now several crystal structures of full length (or nearly full length) rat and human PAH (e.g. Fig 1B) that more thoroughly define the structured and disordered regions of the RS-PAH conformation. In these, the N-terminal ∼20 amino acids, the active site lid, and the last 3-6 C-terminal amino acids are disordered. Differences in subunit positioning among these structures suggests significant mobility within a C-terminal 4-helix bundle, labeled in the right-most image in Fig. 1B (5,11). In addition, there are now a series of publications that address improved modelling of the A-PAH conformation first introduced in 2013, the most recent of which is illustrated in Fig 1C (5,7,11-14). However, no crystal structure exists for full-length PAH (rat or human) in the A-PAH conformation, and little experimental attention has been given to the mechanism for the transition between the RS-PAH and A-PAH conformations.

The transition between RS-PAH and A-PAH conformations minimally involves a dramatic reorientation of protein domains within each protomer of the multimers (compare Figs 1B and 1C). The reorientation has been proposed to involve the N-terminal 25 – 30% of each protomer (residues 1-117 or residues 1-128; segments 1-4) (4,13). Specific to the RS-PAH conformation is an intra-subunit auto-inhibitory interaction at residue ∼23 (in segment 2), which partially occludes the enzyme active site (within segment 6), presumably dictating low activity. The A-PAH conformation does not have the autoinhibitory interaction, but contains an inter-subunit interface involving residues ∼43-77 (within segment 3, a.k.a. the ACT domain), which can be stabilized by allosteric Phe binding at this interface through a conformational selection mechanism (4,15). ACT domains serve as ligand sensors in many proteins (16-18). The A-PAH conformation has a more accessible active site, presumably dictating high activity.

The Phe-modulated equilibrium among alternate PAH conformations allows PAH to respond to protein intake or catabolism and regulate blood Phe with a more nuanced sensitivity than would be possible for K_M_-dependent substrate level control. An individual’s basal Phe level can simplistically be thought of as the equilibrium tipping point between the RS-PAH and A-PAH conformations (3). The tipping point for most humans is in the range of ∼50 – 120 μM Phe, while individuals living with PKU can exhibit significantly higher Phe levels (300 μM – 2.5 mM). PKU is a highly heterogeneous recessive disorder, with more than 1000 disease-associated alleles (19,20). PKU treatment is further complicated by the fact that the neurologic consequences of a given Phe level can vary considerably among individuals. Treatment strategies under development are diverse (21), many of which would benefit from a deeper understanding of the conformational space available to PAH.

The tetrameric RS-PAH conformation is well described for full-length mammalian PAH (Fig 1B), with the exception of the aforementioned disorder (e.g. PDB: 5DEN, 5FGJ, 6N1K) (5,11,13). Alternate models exist for the A-PAH conformation, some of which are supported by small angle X-ray scattering (SAXS) data, and a crystal structure of allosteric Phe bound to a portion of the protein (residues 34-111, segment 3, the ACT domain, PDB: 5FII) (4,5,7,11,13,24). One such SAXS-supported A-PAH conformation model is illustrated (Fig 1C) (13); it places the dimerized regulatory domain considerably farther from the protein’s center of mass relative to our original, other previous, and contemporary models (5,7,11,24).

There is precedent for single amino acid variants that dramatically alter a protein’s multimeric equilibrium (e.g. (25,26)) and reveal the structural basis for an inborn error of metabolism (27). We have proposed that some disease-associated single amino acid variants of PAH can contribute to PKU by shifting the equilibrium between the RS-PAH and A-PAH conformations, thus altering the Phe concentration required to promote activation (i.e. raising the tipping point) (3). To further probe the RS-PAH ⇔ A-PAH transition, herein we characterize variants designed to destabilize the RS-PAH conformation by disrupting a single intra-subunit interaction. The targeted interaction is a cation-π sandwich, which, depending on the environment, has been estimated to introduce between −45 and +30 kcal/mole to protein stability, with stabilization enhanced by neighboring residues that neutralize the repulsion between the two cations (28). Figure 2A illustrates the targeted cation-π interaction between Phe80 in the ACT domain, and Arg123 and Arg420, which are near the N-and C-termini of the catalytic domain (see Fig 1A). In this interaction, charge neutralization is potentially provided by Glu66, with Arg68 likely involved in stabilizing the hydrogen bonding network (5). Substantial domain repositioning in the transition from RS-PAH to A-PAH (see Fig 1C), dictates that this intra-subunit cation-π interaction at Phe80 *cannot* be present in any of the proposed A-PAH conformations (Fig 2B). Although various models for the A-PAH conformation differ in their details, all contain the ACT domain dimer and all place the Phe80 side chain as available for intra-subunit interactions with the catalytic domain and inter-subunit interactions with C-terminal residues.

**Figure 2:**
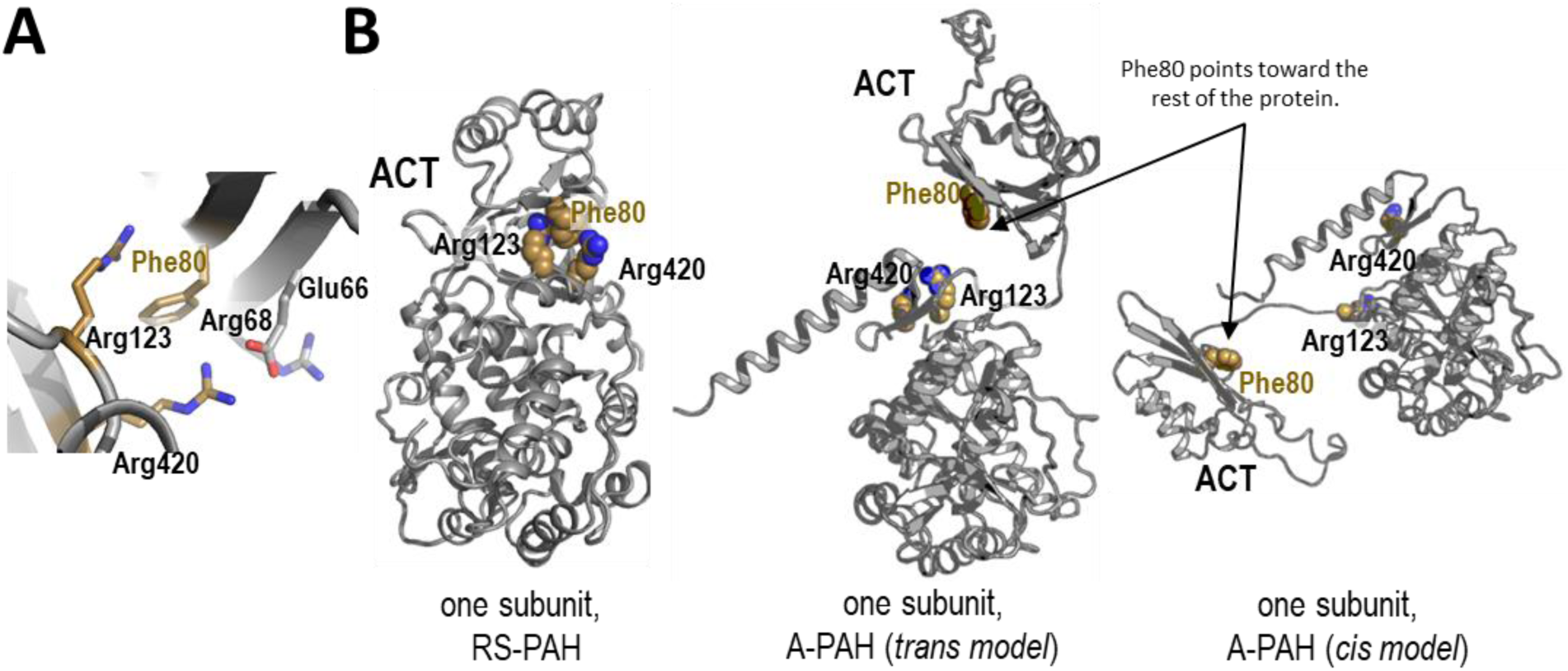
The juxtaposition of Phe80, Arg123, and Arg420 of one PAH protomer in alternate conformations. (A) The cation-π sandwich first identified in the rPAH crystal structure (5), with potential charge neutralization provided by Glu66 and involvement of Arg68. (B) Alternate positioning of residues Phe80, Arg123, and Arg420 (shown in space filling) in different PAH conformations. The RS-PAH crystal structure (left) shows the cation-π sandwich. Both the commonly assumed A-PAH trans conformation (center) and the recently proposed A-PAH cis conformation (right) (13) cannot include cation-π interactions between Phe80, Arg420, and Arg123.

Herein we describe the results of substituting Phe80 with alanine, leucine, arginine, and aspartic acid. These substitutions all remove the RS-PAH-specific cation-π sandwich, which might be predicted to destabilize the RS-PAH conformation and favor formation of the A-PAH conformation. Substitution with alanine largely removes side chain interactions. Substitution with leucine retains the hydrophobic side chain character. Substitution with arginine imposes a charge repulsion in the RS-PAH conformation. Substitution with aspartic acid potentially adds an RS-PAH-stabilizing salt bridge. We cannot yet predict how these substitutions impact the A-PAH conformation, where we do not know the molecular interactions of Phe80, as an atomic resolution structure is lacking. We can however predict, based on all A-PAH models, that Phe80 is expected to extend from the inner surface of the ACT-domain dimer towards the rest of the protein (see Figs 1C and 2B). Herein we present characterization of the F80A, F80D, F80L and F80R variants as a function of Phe concentration using a variety of tools that are established to probe the conformation of mammalian PAH (5,13,15). These include purification characteristics (e.g. resolution of conformers, native polyacrylamide gel electrophoresis (PAGE)), intrinsic protein fluorescence, size exclusion chromatography (SEC), ion exchange chromatography (IEC), enzyme kinetics, and limited proteolysis. The results provide significant insight into PAH conformational dynamics, and point to the power of using multiple complementary approaches.

The PAH conformational landscape includes the characterized RS-PAH conformation, the extensively modeled A-PAH conformation, as well as short-lived intermediate conformations/assemblies that are difficult to isolate or characterize. However, it has long been realized that PAH activation minimally involves releasing autoinhibition through movement of the N-terminal 24 – 30 amino acids (segment 2, which would by necessity drag along segment 1) (29,30). A recent report on a constitutively active rat PAH (rPAH) model of a disease-associated human PAH (hPAH) variant, R68S, suggests the existence of conformations wherein autoinhibition is relieved but ACT domain dimerization is not present, thus uncoupling these phenomena (14). One interpretation of a variety of biochemical and biophysical data is that the R68S variant may be “undocked” such that the entire regulatory domain lacks intra- and inter-subunit interaction with the catalytic domain (as in the RS-PAH conformation, see Fig 1B) nor an intersubunit interaction with a neighboring ACT domain (as in the A-PAH conformation, see Fig 1C), thereby increasing solvent accessibility of loops and protein interfaces that are otherwise protected. The undocked interpretation is consistent with high thermal instability for R68S (14), studies showing that this amino acid substitution interferes with allosteric Phe binding (31), and *in vivo* studies indicating enhanced protein degradation relative to wild type (32,33). In the following description of the Phe80 variants, we find it useful to consider RS-PAH, A-PAH, as well as putatively undocked intermediate conformations. Two examples of putative models of such undocked conformations are illustrated in Fig 1D.

## 2. EXPERIMENTAL PROCEDURES

### 2.1 Protein production and purification methods

Proteins were overexpressed heterologously in an *E. coli* system using a cleavable N-terminal His-SUMO affinity tag as previously described (13,34). Wild type (WT) rPAH was also expressed and purified using the Shiman method as previously described (4,5,10,13). The plasmids encoding the variants were produced by Genewiz. The resultant PAH contains no non-PAH residues at either end of the full-length PAH protein. The classic two-step purification on Ni-Sepharose using a Tris-HCl buffer as previously described (13) yielded nearly homogeneous PAH per SDS PAGE. The final purification step uses IEC fractionation (either a 1-mL or 5-mL HiTrap Q column) in 30 mM Tris-HCl, pH 7.4, 15% glycerol, and a 30-column volume linear gradient to 400 mM KCl. IEC fractionation of SDS-pure rPAH is established to separate quaternary structure isoforms (4,5). Two-mL fractions were collected, analyzed by SDS and native PAGE, flash frozen and stored at −80 °C. The preparative IEC column on Phe80 variants (∼15 – 50 mg per 1 mL column run) yielded 5 – 7 two-ml fractions at ∼0.5 - ∼6 mg/ml giving a final overall yield of ∼ 7 mg purified protein/g cells, for each variant. In some instances, where the HiTrap Q column had been heavily loaded (e.g. >∼25 mg for the 1-mL column), fractions were pooled, diluted with elution buffer to a final salt concentration of <20 mM KCl, and the column was re-run in an attempt to isolate fractions containing only one quaternary structure isoform. PAH is routinely stored (either at 4 °C or −80 °C) as the preparative Hi-Trap Q fractions; once brought to room temperature, PAH is allowed to equilibrate at RT for ∼1 h prior to experimental observations.

### 2.2 Analytical ion exchange chromatography

For each of the PAH proteins, one or more of the preparative IEC column fractions was used for an analytical IEC investigation of the effect of Phe on the separable conformations. For these studies, 100 µg of protein is diluted to 5 ml of 30 mM Tris-HCl, pH 7.4, 20 mM KCl, the sample is pumped onto a 1 ml HiTrap Q column (0.5 ml/min), followed by a 40 min wash of the same buffer containing a set concentration of Phe (0, 10, 30, 100, 300, and 1000 µM). Elution used a 30 ml linear gradient to the same column buffer at 400 mM KCl. Blank gradients (one or more) were performed at each Phe concentration prior to each protein injection.

### 2.3 Intrinsic PAH fluorescence methods

Intrinsic fluorescence of PAH at a concentration of 0.5 μM subunit is measured at 25 °C by exciting the protein solution with a light of wavelength 295 nm and collecting the emission spectrum between 305 and 400 nm. Intrinsic fluorescence measurements were obtained for each of the Phe80 variants and for WT rPAH in the absence and presence of 1 mM Phe.

### 2.4 PAH enzyme kinetics

PAH activity was determined at 25 °C by continuously monitoring tyrosine fluorescence intensity after initiating the assay by addition of PAH, which was at a concentration between 0.1 and 0.4 mg/mL (i.e. ∼ 2-8 μM subunit). Prior to addition to the assay mixture, the protein was preincubated with or without Phe, at the assay concentration, for 1 minute at room temperature. The continuous fluorescence-based assay is based on that of Gersting *et al*. (35) with modifications introduced by Bublitz (36). Tyrosine fluorescence measurements use an excitation wavelength of 275 nm and an emission wavelength of 305 nm on a PTI Fluorescence System spectrophotometer equipped with an USHIO Xenon Short Arc Lamp and Felix™ for Windows Software. The standard reaction is performed in a final volume of 2 mL containing 20 mM bis-tris propane (pH 7.3), Phe (at concentrations between 10 µM and 1 mM, Fig S3), 10 µM ammonium iron (II) sulfate, 40 µg/mL catalase (800 – 3200 units), 75 µM BH_4_ and 75 µM DTT. Initial velocity reflects the first 50 sec. The initial velocity data were fitted to a 3-parameter Hill equation using the SigmaPlot v. 10.0 (Systat Software, San Jose, CA). Calibration curves of standard tyrosine concentrations (10, 20, 30, 40 µM in the assay solution) were obtained on the day of the activity analysis.

### 2.5 Native proteolytic digestion

WT rPAH and the Phe80 variants were subjected to native trypsinolysis at 25 °C, at 1 mg/ml and various concentrations of Phe (0, 30 μM, 100 μM, 1 mM), 30mM Tris-HCl pH 7.4, and 150 mM KCl using a PAH:trypsin ratio of 200:1. The digestions were monitored for 1 hour at RT, taking samples at 1, 15, 30, 45, and 60 min; stopping the reaction by addition of 100 μM PMSF. AspN protease digestions used a 1:250 protease:PAH ratio and were monitored at various concentrations of Phe (0, 100 μM, 1 mM). AspN digestion was stopped by exposure to 100 °C for 5 min. Proteolytic digestion was monitored by SDS PAGE using 12.5% PhastGels. For selected digests (WT rPAH and F80R) peptides were identified by submission to the Wistar Proteomics Facility, which used denaturing trypsin digestion and mass spectroscopy. For those examples, the experiments were repeated and BioRad gels were used for peptide separation by SDS PAGE using a range of sample loadings such that bands could be excised for analysis. For WT rPAH, without Phe, trypsinolysis at a 250:1 ratio, was followed for the effect on quaternary structure using native PhastGels alongside the SDS analysis.

### 2.6 Analytical SEC

SEC analyses were performed using a calibrated Superdex 200 10/300 GL column (GE Healthcare) in 30 mM Tris-HCl, pH 7.4, 150 mM KCl, without or with 1mM Phe at a flow rate of 0.5 ml/min at RT. Samples were from individual preparative IEC fractions, containing 15% glycerol, flash frozen, stored at −80 °C, defrosted at 25 °C, and allowed to sit for an hour. Samples were adjusted to a protein concentration of ∼ 1.5 mg/ml for injection onto the pre-equilibrated column.

## 3. RESULTS

### 3.1 Resolution of conformational isoforms for rPAH and the Phe80 variants

Whether purified using the classic phenyl Sepharose affinity method (10), or cleaved from the N-terminal 6-His-SUMO construct, the resulting SDS-pure rPAH (or Phe80 variants) could be resolved into conformationally distinct isoforms using preparative IEC (4). The IEC fractionation profiles provide novel information on the different multimerization properties of Phe80 variants. As reported previously for WT rPAH, SDS-pure PAH fractionates on IEC into distinct conformations as determined by native PAGE (and confirmed by SEC and/or SEC-SAXS) (5). The order of elution with increasing ionic strength is dimer, a faster migrating tetramer, and a slower migrating tetramer. Nearly all of our published work on the RS-PAH conformation of rPAH has used fractions containing predominantly the faster migrating tetramer (4,5,15). Addition of 1 mM Phe to a sample comprised predominantly of a faster migrating tetramer results in loss of dimer (if present), migration to a position similar (on both IEC and native PAGE) to the slower migrating tetramer, and accumulation of higher order multimers (4). Although the structural similarities/differences between different rPAH tetramers observed in the absence of Phe remains unclear, IEC effectively discriminates the behaviors of the Phe80 variants relative to WT rPAH. Below we present native PAGE and analytical IEC as a method to compare preparative IEC fractions from the final purification step of each of the Phe80 variants. A later section presents the use of analytical IEC to explore how the multimeric equilibrium of each of the Phe80 variants responds to the presence of different concentrations of Phe in a physiologically relevant range.

### 3.2 IEC behavior of Phe80 variants in the absence of Phe analyzed by native PAGE and SEC

The preparative IEC behavior of the studied proteins fell into two categories. The first, including WT rPAH, F80A, and F80D, are resolved to show dimer, and two different tetramer conformations as confirmed by native PAGE (Fig 3A). The tetramers appear less well resolved for the F80A and F80D variants relative to WT rPAH. The second category, comprised of F80L and F80R, show a single tetrameric band with little to no evidence for dimer in the preparative IEC fractions (Fig 3B). All variants show evidence of higher order multimers that elute at the higher ionic strength end of the peak. Western blots (not shown) demonstrate these higher order forms to be PAH. The proportion of higher order multimers was variable among preparations. A remarkable property of WT rPAH is the stable composition of the preparative IEC fractions wherein the multimeric components do not re-equilibrate upon lengthy storage at either 4 °C or −80 °C. This implies a high activation energy to the multimeric interconversion process. The same stability could not be demonstrated for F80A or F80D. The second category of variants comprised of F80L and F80R show an (apparent) single tetrameric conformation in the preparative IEC fractions, consistent with the hypothesis that these variants may be in the A-PAH conformation. It is well established that activated PAH is less prone to tetramer dissociation (e.g. (13)), presumably due to the presence of additional/different inter-subunit interactions (e.g. the ACT domain dimer interface). To more fully characterize the conformational equilibrium of WT rPAH and the Phe80 variants in the absence of Phe, analytical IEC was carried out for one or more fractions that had eluted from the preparative IEC columns and analyzed by native PAGE (Fig S2). Taken together with the SEC data in Fig 3, these experiments confirm that the order of elution from the IEC column in the absence of Phe is dimer, faster migrating tetramer, and slower migrating tetramer.

**Figure 3.**
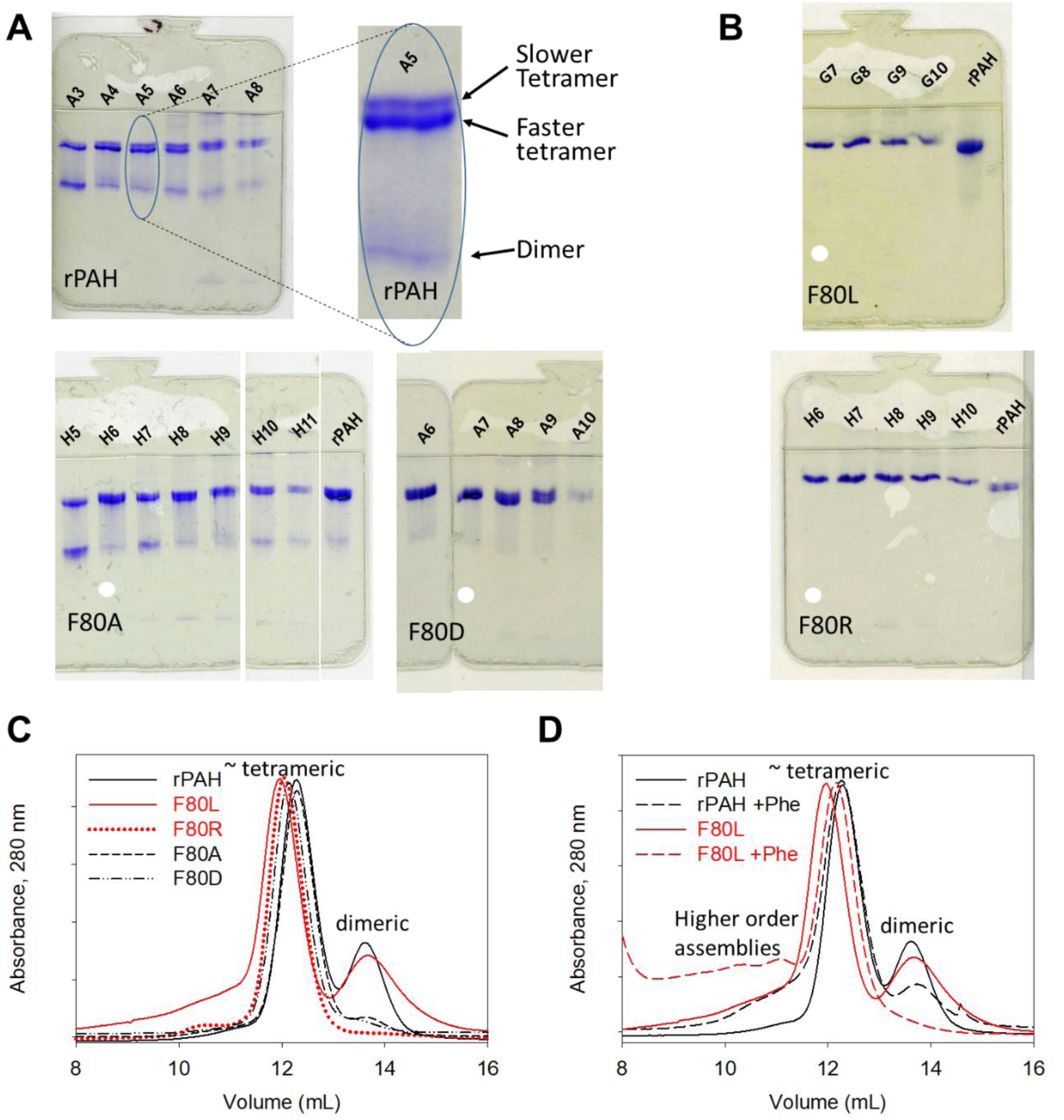
Individual fractions from the preparative IEC purification step were analyzed using 12.5% native PhastGels and analytical SEC. The white dots indicate fractions used for determination of enzyme kinetic parameters. The letter used with fraction numbers was varied among preparations. **(A)** WT rPAH, F80A, and F80 D show multiple quaternary structure isoforms. WT rPAH fractionates into a dimeric and two tetrameric components. Shown right is a 3.5x enlargement of fraction A5. F80A and F80D behave similarly, though the tetramer bands are less sharply resolved. **(B)** Preparative IEC fractions of F80L and F80R show only one tetramer band and no significant dimer. **(C)** Analytical SEC on preparative IEC fractions of WT rPAH and all the Phe80 variants following storage at −80. **(D)** WT rPAH and F80L chromatograms are compared in the presence (dashed lines) and absence (solid lines) of 1 mM Phe in the column buffer.

Some of the IEC fractions illustrated in Figs 3A and 3B were also subjected to analytical SEC (see Figs 3C and 3D), where the elution volumes of tetramer and dimer were previously established (4,13). An SEC comparison of WT rPAH and the Phe 80 variants (Fig 3C) shows variations in elution times, suggesting differences in tetramer dimensions, with F80L being largest, followed by F80R, then F80D, and F80A, which elutes at the same time as WT rPAH. We posit that the apparent average sizes may reflect contributions from the different conformations, with contributions from the A-PAH or undocked conformations causing an apparently larger size. The selected chromatographic fractions also have different dimeric content. The WT rPAH fraction was specifically chosen to contain dimer. However, for the F80L sample, the high dimer content appears to be a result of the freeze/storage/thaw process (compare with Fig S2C). For WT rPAH and F80L, Fig 3D shows the effect of 1 mM Phe on the SEC profile. As we had previously demonstrated, addition of 1 mM Phe in the SEC column buffer causes the WT rPAH peak to elute slightly earlier, consistent with a larger size (4) as predicted by our newest model for the A-PAH conformation (Fig 1C) (13). Addition of Phe also results in a diminished population of WT rPAH dimer and slight accumulation of larger components. For F80L, addition of Phe causes the tetrameric component to convert to the same size as WT rPAH. This is consistent with F80L containing a considerable population of undocked conformations, which can be drawn to the A-PAH conformation through interaction with Phe. Consistent with this, addition of Phe also causes the F80L variant to lose nearly all of its dimer content. Notably, in the presence of Phe, F80L gains in population of higher order multimers. We have posited that the appearance of such multimers are Phe-stabilized species wherein PAH tetramers are tethered together by formation of inter-assembly ACT domain dimers (4). Formation of such inter-tetramer interactions could be facilitated by the undocking ACT domains, leaving them free to interact with ACT domains of nearby tetramers.

### 3.3 PAH intrinsic protein fluorescence as a measure of PAH conformational change

A dramatic change in the PAH intrinsic protein fluorescence upon addition of Phe was one of the first indications that an activated form of PAH had a significantly different conformation relative to its resting state (37). A red-shift of ∼ 10 - 15 nm in the tryptophan emission profile is routinely observed for mammalian PAH following addition of 1 mM Phe, which is sufficient to fully activate WT PAH. This change, illustrated in Fig 4A, has been used both as an assay for PAH conformational change and to better understand the structural basis of PAH activation (37-41). Intrinsic fluorescence is a readout of the environment of the three tryptophan residues at positions 120, 187, and 326. Trp120 (noted in Figs 1B-D) is in the linker region (segment 4) between the ACT domain and the catalytic domain; its environment will change in the RS-PAH ⇔ A-PAH conformational transition (3,4). A detailed study using human PAH concluded that Trp120 contributes ∼ 61% to the total fluorescence emission changes that are Phe-stabilized, while Trp326 and Trp187 (both in segment 6) are less influential (at ∼26% and ∼13%, respectively) (39). The WT rPAH intrinsic fluorescence in the absence of Phe reflects tryptophan residues that are largely buried in nonpolar environments. In the presence of Phe, the spectra reflect one or more tryptophan residues moving to more polar environment, with at least one becoming largely solvent accessible (fluorescence at 350 nm). In addition to illustrating the intrinsic fluorescence of WT rPAH (+/-1 mM Phe), Fig 4A shows the intrinsic fluorescence of the Phe80 variants in the absence of Phe; there is a remarkable similarity in the spectra of F80L and F80R to that of WT rPAH plus 1 mM Phe. In the absence of Phe, the spectra of F80D and F80A look similar to WT rPAH. Addition of 1 mM Phe (Fig 4B) does not dramatically alter the fluorescence spectra of F80L and F80R, but changes the spectra of F80A and F80D to match that of WT rPAH with 1 mM Phe.

**Figure 4.**
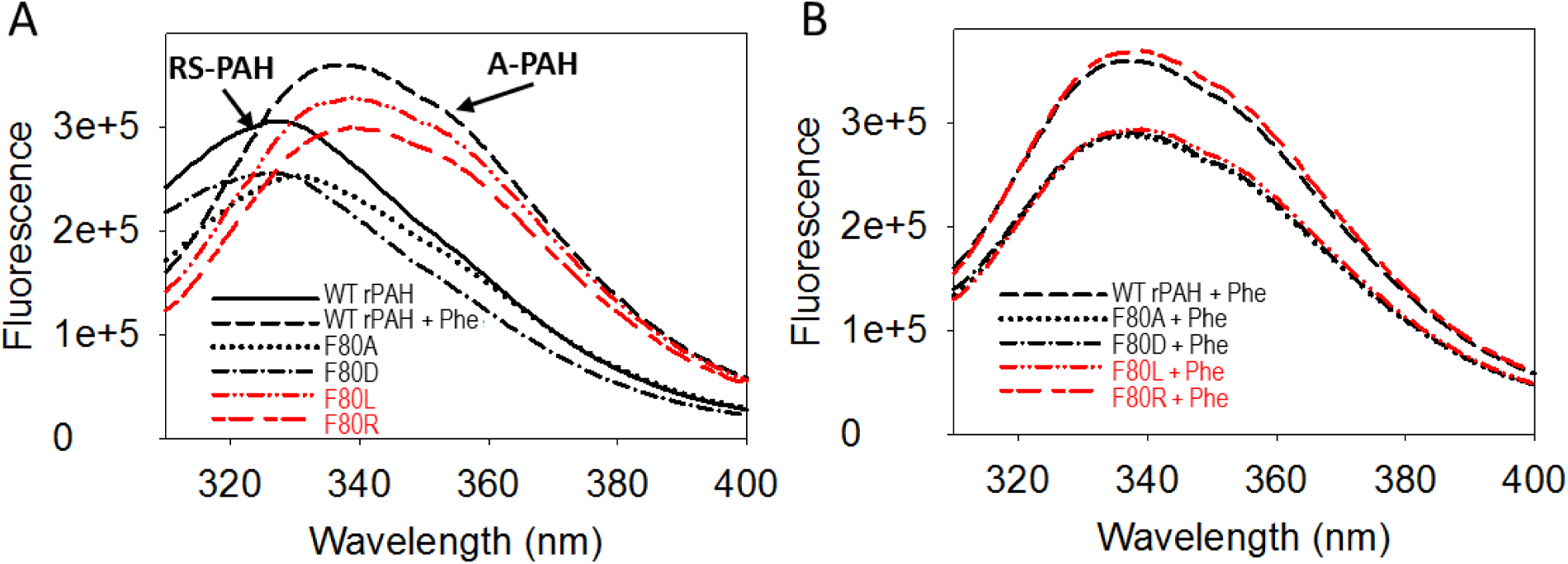
Comparative intrinsic fluorescence of rPAH and the Phe80 variants. **(A)** Emission spectra are shown for rPAH and the Phe80 variants in the absence of Phe. The dashed black line shows WT rPAH in the presence of 1 mM Phe. **(B)** Emission spectra are shown for rPAH and the Phe80 variants in the presence of 1 mM Phe.

The intrinsic fluorescence results initially suggest that F80L and F80R are in an A-PAH conformation while F80A and F80D are in an RS-PAH conformation. We might also conclude that addition of Phe allows each of the Phe80 variants to achieve the A-PAH conformation, as for WT rPAH. Note however that each of these interpretations of the fluorescence data reflects the assumption that the observed red shift reports on a *gain* of the A-PAH population. If, instead, the red shift reports on a *loss* of the RS-PAH population, these results are also consistent with the possibility that the Phe80 variants populate conformations intermediate between RS-PAH and A-PAH, where Trp120 is solvent accessible. The suggested conformations have the ACT domain undocked from the intra- and inter-subunit interactions shown for both the RS-PAH or A-PAH conformations (labelled in Figs 1B and 1C); two modelled examples of undocked conformations are illustrated in Fig 1D. Thus, an equally rational interpretation of the intrinsic fluorescence data is that loss of the cation-π interaction at Phe80 increases the propensity of PAH to sample undocked conformations.

### 3.4 Enzyme Kinetics

For determination of apparent K_M_ and V_MAX_ values for WT rPAH and Phe80 variants, we capitalize on the sensitivity of PAH activity to pre-incubation with Phe simplistically as a proxy for the position of the RS-PAH ⇔ A-PAH conformational equilibrium. For example, the initial velocity of WT rPAH is higher when the enzyme is incubated with Phe prior to starting the assay relative to when the enzyme is not preincubated with Phe. This phenomenon arises from preincubation with Phe as stabilizing the A-PAH conformation and increasing its population. For the Phe80 variants, kinetic parameters were determined for one fraction off the preparative IEC column, as noted in Fig 3. Table 1 includes the fitted kinetic values; Figure 5 shows the fits of the experimental data. A classic approach to evaluating PAH variants is to quantify the fold activation observed when the protein is preincubated with Phe relative to no preincubation. Here we see the expected ∼5-fold activation (reported as comparison of V_MAX_) for WT rPAH. F80D and F80A show lower fold activation of 3.2 and 1.8 respectively, consistent with the notion that these amino acid substitutions destabilize the RS-PAH conformation, making it easier to attain the A-PAH conformation upon addition of Phe. F80L and F80R are constitutively active, showing fold activations of ∼1. Although this is consistent with F80L and F80R as constitutively in the A-PAH conformation, a closer look at the kinetic parameters for all the Phe80 variants can also be interpreted as supporting constitutively active undocked conformations. Like the A-PAH conformation, undocked conformations are expected to have fully accessible enzyme active sites.

**Table 1:** Kinetic Parameters for WT rPAH and Phe80 variant in the absence or presence of a 1 min preincubation with Phe (at the assay concentration).

| PAH variant | $K_M$ ($\mu M$) | $V_{MAX}$<br>(nmol Tyr/min/mg) | Hill<br>coefficient | Fold<br>Activation | V/K |
| --- | --- | --- | --- | --- | --- |
| WT rPAH (-Phe) | 203 ± 45 | 465 ± 46 | 2.2 ± 0.8 | | 2.2 |
| F80D (-Phe) | 327 ± 107 | 1,210 ± 230 | 1.6 ± 0.5 | | 3.7 |
| F80A (-Phe) | 229 ± 18 | 3,470 ± 141 | 1.8 ± 0.2 | | 15.1 |
| F80L (-Phe) | 59 ± 3 | 7,600 ± 147 | 1.3 ± 0.1 | | 129.0 |
| F80R (-Phe) | 78 ± 8 | 4,100 ± 151 | 1.6 ± 0.3 | | 52.6 |
| WT rPAH (+Phe) | 158 ± 58 | 2,560 ± 337 | 2.6 ± 1.7 | 5.5 | 16.2 |
| F80D (+Phe) | 142 ± 12 | 3,890 ± 164 | 3.2 ± 0.7 | 3.2 | 27.4 |
| F80A (+Phe) | 180 ± 15 | 6,310 ± 208 | 2.2 ± 0.2 | 1.8 | 35.1 |
| F80L (+Phe) | 65 ± 6 | 8,130 ± 282 | 1.2 ± 0.1 | 1.1 | 125.0 |
| F80R (+Phe) | 79 ± 7 | 3,880 ± 142 | 1.4 ± 0.2 | 0.9 | 49.1 |

**Figure 5.**
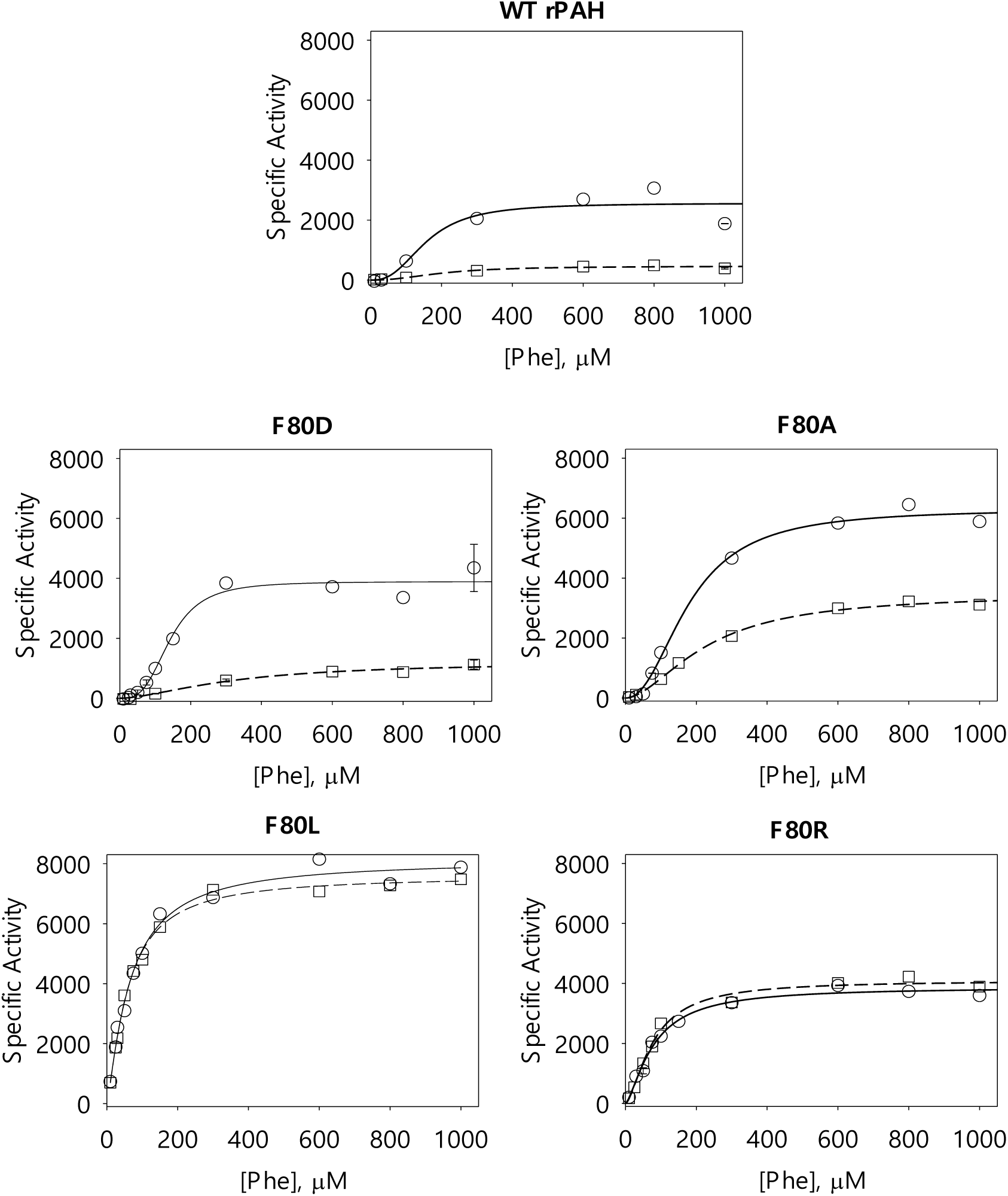
Kinetic data for WT rPAH and Phe80 variants fit to a three parameter Hill equation. Specific activity is reported as nmol Tyr min^-1^ mg^-1^ (see Fig 3 for identification of selected IEC fractions). Assays performed with and without Phe preincubation are shown in circles and squares respectively with regression lines shown as solid and dashed respectively.

The K_M_ values for PAH reflect a complex function of Phe binding both to the active site and to the allosteric site at the ACT domain dimer interface. Table 1 shows that the K_M_ values, like the native PAGE and SEC data, fall into two groups. For WT rPAH, F80D, and F80A, the K_M_ values are all within experimental error of one another, both without preincubation (∼250 μM) and with preincubation (∼155 μM). In contrast, the K_M_ values for F80L and F80R are significantly lower (∼70 μM) and insensitive to the preincubation step. We also note that all of the Phe80 variants show V_MAX_ values larger than WT rPAH (under comparable conditions). One way to compare such values is to calculate a simple V/K ratio. Because the kinetics of PAH is a function of a complex allosteric activation by substrate, the numerical values of the V/K ratio are not mechanistically meaningful. However, Table 1 shows a dramatic variation in V/K suggesting that the rate-limiting step for WT rPAH, which is likely a conformational change, is relieved in the Phe80 variants. The expected conformational flexibility of undocked PAH is consistent with relief of a rate limiting conformational change, perhaps involving breaking the Phe80 cation-π sandwich. Finally, we note that the Hill coefficients for WT rPAH, F80D, and F80A are not significantly different from one another (∼2), and likely reflect the formation of two ACT domain dimers per activated tetramer. Our published computational analysis of Phe binding to an isolated ACT domain dimer established the independent (non-cooperative) binding of two Phe molecules (15); thus binding of one Phe to each ACT domain dimer of a tetramer will afford full activation.

### 3.5 Analytical IEC as a function of Phe

We had previously established that addition of 1 mM Phe to WT rPAH results in a change in an analytical IEC profile using 100 μg protein on a 1 ml HiTrap Q column, and that ∼50 μM Phe shows a profile containing peaks consistent with both the RS-PAH conformation (no Phe) and the A-PAH conformation (1 mM Phe) (4). Herein we use a slightly different procedure (see Fig 6) to compare and contrast the conformational sensitivity of WT rPAH and the Phe80 variants to Phe. This study provides additional evidence that all the Phe80 variants are neither fully RS-PAH-like nor fully A-PAH-like in the absence of Phe, notably Fig 6A shows that the IEC chromatographic behavior of all the studied proteins responds to the addition of Phe. Fig 6B shows the progressive change for WT rPAH as Phe is varied from 0 to 1 mM in ∼half-log increments. The chromatograms reflect a decrease in the amount of dimer present (first notable at 30 μM Phe), a transition to the A-PAH conformation (the dominant species at 100 μM Phe), and accumulation of higher order multimers (significant by 300 μM Phe). For the F80A and F80D proteins, shown respectively in Fig 6C and 6D, the chromatograms show similar effects, with the exception that intermediate Phe concentrations (e.g. 30 μM) show the mixture of isoforms less well resolved. For WT rPAH, F80A, and F80D, the predominant peak in the absence of Phe elutes at 30.8, 31.0, and 31.2 min, respectively, while the predominant peak at 100 μM Phe elutes at 32.0, 31.8, and 32.0 min respectively (see Fig 6A). In the absence of Phe, F80L and F80R show single peaks, whose elution times are at intermediate values, 31.6 and 31.3 min respectively (see Fig 6A). Addition of Phe alters these elution times to 31.9 and 31.8 min respectively. Notably for both F80L (Fig 6E) and F80R (Fig 6F), this effect is seen at Phe concentrations significantly lower than their K_M_ values, which are ∼70 μM, and likely reflect Phe binding to the allosteric site on the ACT domain dimer. For F80L, the effect saturates at 10 μM Phe; by 30 μM Phe F80L shows about 50% formation of other assemblies (eluting at 32.8 min). Independent data show that the other assemblies are likely higher order multimers (see Fig 3D). In contrast, for F80R, the change in elution time saturates at 30 μM Phe and there is little evidence for accumulation of alternate assemblies at 100 μM Phe. From these results, we feel confident in concluding that F80L and F80R are not fully A-PAH-like in the absence of Phe, in support of the hypothesis that Phe80 side chain interactions are important to both the RS-PAH and A-PAH conformations.

**Figure 6:**
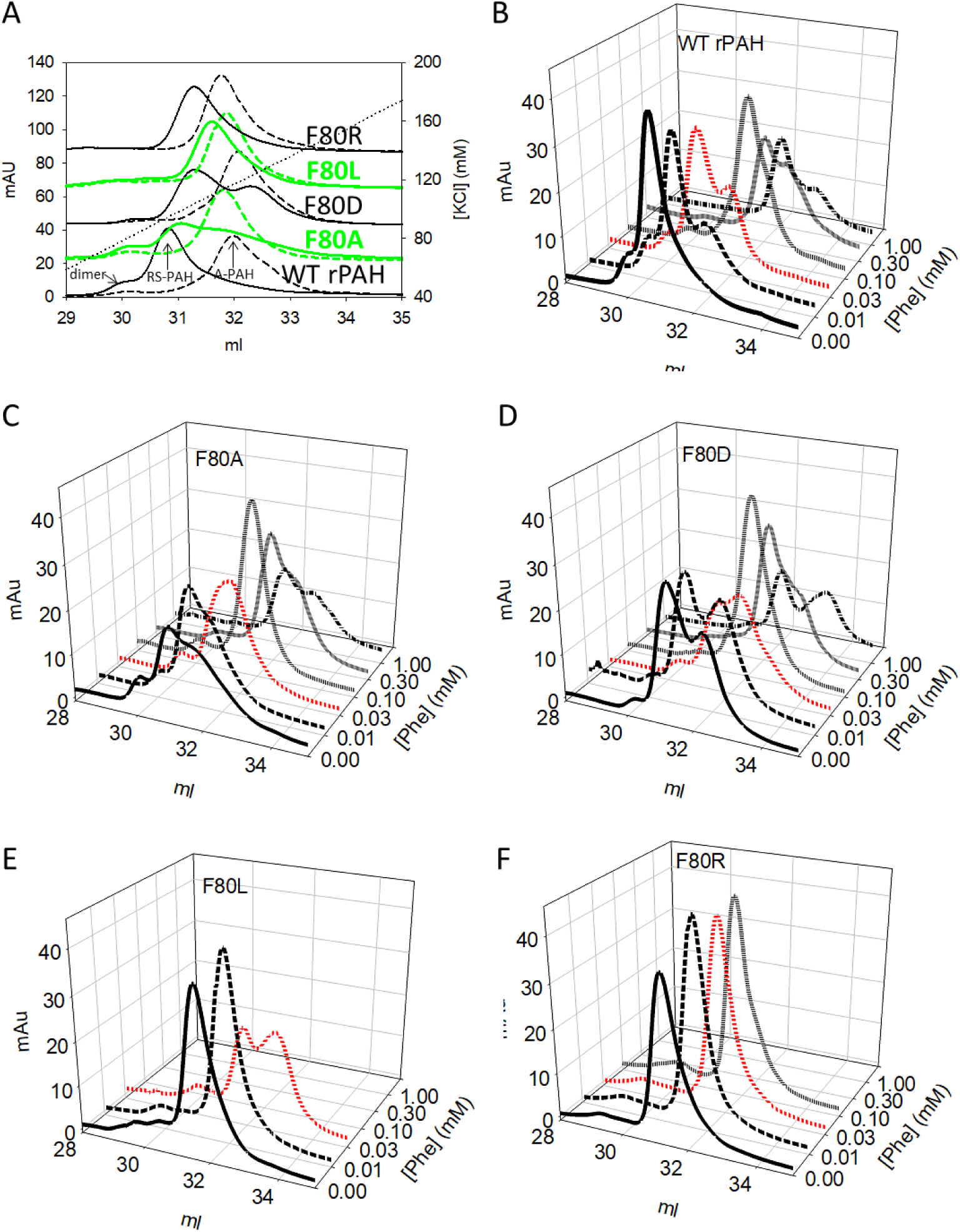
Analytical IEC on rPAH and Phe80 variants as a function of [Phe]. For all separations, 100 μg of protein was diluted into 5 ml start buffer (no Phe) and applied to a 1 ml HiTrap Q column that had been pre-equilibrated with the set concentration of Phe (10 min). The column was washed with is starting buffer for 40 min and protein isoforms were eluted with a linear gradient to 400 mM KCl at the set Phe concentration. Part A provides a comparison of peak positions in the absence of Phe and at the Phe concentrations that corresponds to the predominant peak in the A-PAH conformation. Alternating black/green coloring is used to discriminate among partially overlapping chromatograms. For WT rPAH, F80A, and F80D, this is 100 μM Phe; for F80L it is 10 μM Phe; for F80R it is 30 μM Phe. Parts B-F illustrate the progression of chromatographic changes for each variant as the Phe concentration is increased. To facilitate comparison, the chromatogram at 30 μM Phe is shown in red.

### 3.6 Limited Proteolysis

Limited proteolysis is a classic approach to monitoring conformational change and was first applied to PAH by S. Kaufman and coworkers in the 1980s (42). Herein, we probe the conformations of WT rPAH and the Phe80 variants using trypsin (Fig 7) and AspN protease (Fig S3), monitoring limited proteolysis under native conditions at various concentrations of Phe for a 1 h time course at 15-minute intervals. The results provide the strongest evidence that the Phe80 variants sample conformations that are different from either RS-PAH or A-PAH and are highly susceptible to proteolysis in the linker region between the regulatory and catalytic domains (segment 4), as would be predicted for undocked conformations. In addition, proteolysis reveals facile hydrolysis within the C-terminal 4-helix bundle when PAH is in the RS-PAH conformation, consistent with transient tetramer dissociation. Proteolytic cleavage sites are mapped onto the protomer structures shown in Fig 1 and included in the sequence alignment illustrated in Fig S1.

**Figure 7:**
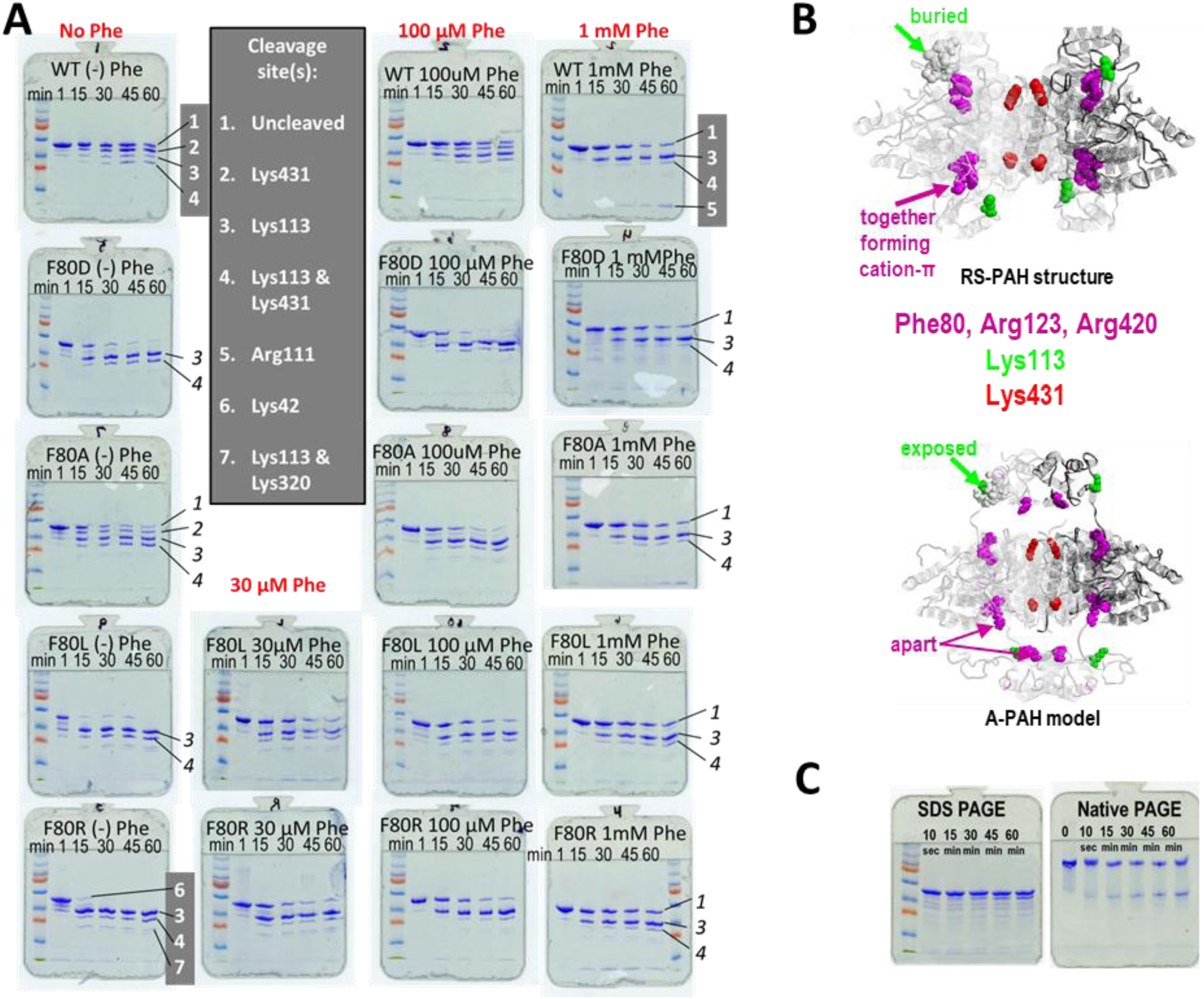
Tryptic digests as a function of time and [Phe]. **(A)** SDS PhastGel analysis of a native tryptic digest time course for WT rPAH and the Phe80 variants. Each gel is entitled with the protein name. Each column is topped with the concentration of Phe present. The lanes are labeled with the reaction times of 1min, 15 min, 30 min, 45 min, and 60 min. The leftmost lane contains pre-stained molecular weight standards. Peptides were identified for WT rPAH (+/-Phe) and F80R (-Phe) by mass spectroscopy in the Wistar Proteomics Facility on matched samples run on standard mini-gels (numbered bands, white on grey). For all other bands numbered in italics, identification is by inference. A similar study using AspN protease is shown in Fig S3. The tryptic cleavage sites are noted in structures in Fig 1B and 1C; cleavage sites for both proteases are in the sequence alignment in Fig S1. **(B)** Images of the RS-PAH structure and the A-PAH model illustrating the positions of the residues that participate in the cation-π sandwich (purple), as well as two major cleavage sites in green and red. **(C)** Trypsinolysis of WT rPAH in the absence of Phe was monitored in parallel by SDS and native PhastGel analysis. These data show that the accumulation of dimer parallels the appearance of cleavage at Lys431.

Native trypsinolysis of WT rPAH and the Phe80 variants is detailed by SDS PAGE in Fig 7A. The top row of SDS PhastGels illustrates the proteolytic time course for WT rPAH at 0, 100 µM and 1 mM Phe. In this case, the most susceptible site in the absence of Phe, known to reflect the RS-PAH conformation, is Lys431 (see Figs 1B, 7B), whose cleavage can be seen to slowly increase throughout the time course. Lys431 is within the C-terminal helix (residues 426-453) and predicted to be largely inaccessible in the tetrameric RS-PAH crystal structures (e.g. PDB: 5DEN). Native proteolysis within this C-terminal 4-helix bundle provides extremely strong evidence that tetramer dissociation (to dimers or monomers) is facile in the resting state. Cleavage within the 4-helix bundle is predicted to prevent tetramer reformation. This is further illustrated in Fig 7C, which uses matched samples on SDS and native PAGE to show that the accumulation of dimer with time parallels cleavage at Lys431 (using a PAH:trypsin ratio of 1:250). In contrast, at 1 mM Phe, where WT rPAH is in the A-PAH conformation, Lys113 is the primary cleavage site (position noted in Fig 1C and 7B), and the time course shows a more rapid cleavage. Lys113 is within a highly basic region identified to respond to Phe with increased proteolytic susceptibility in Kaufman’s original studies (43). That Phe-stabilized activation results in rapid cleavage at Lys113 is consistent with an extended structure for segment 4 (see Fig 1A) in the A-PAH conformation (Fig 1C and 7B, bottom). In the RS-PAH conformation Lys113, though on the protein surface, is involved in intra-subunit interactions with the carboxyl groups of Asp315 and Asp27, tethering segments 2, 4, and 6 (see Fig 1A). Because the equilibrium of PAH conformations is dynamic, it is not surprising that in the absence of Phe we also observe minor products indicating cleavage at Lys113 as well as products showing cleavage at both Lys113 and Lys431. Likewise, in the presence of Phe, discussed below, we observe minor products indicating cleavage at Lys431. The additional N-terminal peptide identified as cleavage at Arg111 for WT rPAH at 1 mM Phe is due to the RDK sequence from 111-113.

The right most column in Fig 7 shows that all four Phe80 variants show trypsinolysis patterns at 1 mM Phe that are qualitatively indistinguishable from WT rPAH, consistent with the intrinsic fluorescence data and with the hypothesis that all can reach the A-PAH conformation in the presence of sufficient Phe. However, the left most column of Fig 7 shows that all of the Phe80 variants in the absence of Phe show proteolytic patterns that are unlike WT rPAH in both the absence and the presence of 1 mM Phe. This is the strongest evidence that the variants are not in either the RS-PAH conformation nor the A-PAH conformation. In the absence of Phe, Lys431 *is not* a primary cleavage site for any of the Phe80 variants; though it is evident as a minor site in the F80A sample. From this, we might conclude that the F80A variant is most like WT rPAH in its ability to populate the RS-PAH conformation (see also Fig 4A). Instead, all of the Phe80 variants are cleaved at Lys113 in the absence of Phe, and this cleavage occurs much more rapidly relative to WT rPAH in the presence of Phe. This hypersensitivity of cleavage at Lys113 is also consistent with the Phe80 variants sampling conformations that are neither RS-PAH nor A-PAH, as would be expected for undocked conformations. In all the Phe80 variants, addition of Phe *protects* the hypersensitivity of Lys113 to tryptic cleavage, presumably by drawing the proteins into a Phe-stabilized A-PAH conformation. To titrate the sensitivity of the Phe80 variants to protection at Lys113, native trypsinolysis was carried out at intermediate Phe concentrations, as shown in the center two columns of Fig 7. For F80D, F80A, F80L, and F80R protection becomes apparent at > 100 μM Phe, ∼100 μM Phe, 30 μM Phe, and < 30 μM Phe, respectively. These Phe concentrations are consistent with the analytical IEC data (Fig 6) and further supports the hypothesis that all of the Phe80 variants, in the absence of Phe, can sample an open conformation wherein segment 4 (the linker between the ACT domain and the catalytic domain) is hyper-susceptible to trypsinolysis, consistent with undocked conformations. The F80R variant, in the absence of Phe, reveals two additional cleavage sites by the appearance of minor products; these are Lys42 and Lys320. Lys42 is at the edge of a mobile loop that allows the allosteric Phe to bind to the ACT domain dimer interface in the A-PAH conformation (15). In the RS-PAH conformation, its neighbor Glu43 is buried and hydrogen bonded to Tyr204 and His208 of a neighboring subunit. This suggests that F80R contains some population that contains the ACT domain dimer. We cannot comment on the susceptibility of Lys320 at this point.

Native digestion with AspN protease (Fig S3) shows SDS gel patterns that are fully consistent with the conclusions drawn from the trypsinolysis studies (Fig 7). Although the AspN protease generated peptides were not identified by mass spectroscopy, side-by-side comparison of the cleavage patterns suggests that WT rPAH without Phe is susceptible to cleavage at Asp425 and Asp435, again suggesting considerably instability in the 4-helix bundle that secures the RS-PAH tetramer. In the presence of 1 mM Phe, AspN protease appears to cleave WT rPAH at Asp112, consistent with the exposed linker region (segment 4 in Fig 1A) as expected for the A-PAH conformation. Again, in the absence of Phe, all of the Phe80 variants are more susceptible to cleavage in this linker region, showing hypersensitivity (least so for F80A) that can be protected by addition of Phe. With the exception of F80D, where peptides resulting from AspN protease were not identified, all variants achieve a cleavage pattern expected for the A-PAH conformation at 100 μM Phe. Nevertheless, F80D achieves the expected A-PAH-like cleavage pattern at 1 mM Phe.

The limited proteolysis data uniformly support the hypothesis that substitution at Phe80 destabilizes both the RS-PAH conformation and the A-PAH conformation and promotes population of intermediate states for which the regulatory domain is undocked (possible examples in Fig 1D). Prior limited trypsinolysis studies uniformly focused on differential cleavage within a “hinge region” between the regulatory and catalytic domains (residues 111-117) (e.g. (14,44,45)). Interestingly, we could not identify a previous report of cleavage within the C-terminal helices (Lys431 in Fig 7; Asp 425 & 435 in Fig S3), despite one study wherein this cleavage is present in the illustrated SDS gels (14). Here we report prominent C-terminal helix cleavage for WT rPAH in the absence of Phe, which is protected in the presence of Phe. This observation is consistent with many reports that Phe stabilizes tetrameric PAH (e.g. (4)). Nevertheless, the observation of proteolysis within the C-terminal helices, which should be a fully protected 4-helix bundle in the tetramer, argues that tetramer dissociation is facile and raises the question as to the physiologic relevance of dimers or monomers in PAH allostery. Our original proposal that activation of PAH was accompanied by ACT domain dimerization also proposed that tetramer dissociation might be part of the allosteric mechanism, consistent with the morpheein model of protein allostery (4).

Although the mechanistic significance of monomers or dimers in PAH allostery remains unclear, tetramer dissociation and reassociation can contribute to interallelic complementation seen in PKU patients, the majority of whom are compound heterozygotes (e.g. (46,47)).

## 4. DISCUSSION

### 4.1 Phe80 variants of PAH

Herein we report the effects of altering a cation-π sandwich that we had identified as stabilizing the RS-PAH conformation (WT rPAH, PDB: 5DEN) (5); this remains the highest resolution of all structures determined to date for full length mammalian PAH proteins. Phe80, the aromatic component of this interaction was replaced by alanine, leucine, aspartic acid, and arginine. All of the Phe80 variants expressed as soluble proteins, could be isolated in large quantities, stored stably, used for extensive biochemical characterizations, which took place over many months, and all could assume an A-PAH conformation upon addition of Phe. Characterization of the Phe80 variants did not reveal grossly aberrant behavior, with the exception that F80L showed an enhanced propensity to reversibly form discrete higher order multimers. Some of the reported characterizations, if taken in isolation, could be simplistically interpreted as indicating that F80A and F80D are RS-PAH-like in the absence of Phe, while F80L and F80R are A-PAH-like under these conditions. However, the sum of many different types of analyses, most notably the limited proteolysis time course analysis, show that all of the Phe80 variants, to some extent, also sample conformations that are neither RS-PAH-like nor A-PAH-like, but rather more likely reflect undocked intermediates in the allosteric regulation of PAH.

The undocked conformations of the Phe80 variants are presumed on-pathway in the RS-PAH ⇔ A-PAH interconversion for several reasons. First, the proteins are all highly soluble, quite stable, highly active, and thus unlikely to be misfolded. Second, all the proteins can be drawn into the A-PAH conformation through addition of low (and physiologically relevant) concentrations of Phe. Third, in the transition from a conformation wherein the ACT domain is docked to the catalytic domain of one neighboring subunit to a distant location wherein the ACT domain is docked to the ACT domain of a different neighboring subunit must involve a pathway during which the ACT domain is not docked to either neighboring subunit. Like the reported behavior of human porphobilinogen synthase, this is another example where single amino acid substitutions dramatically alter the conformational space sampled by a medically relevant protein and unmask structural conformers that are, in the wild type protein, too short-lived or in low abundance for extensive characterization (48,49). These variants can also provide a glimpse into the aberrant conformational sampling of disease-associated variants.

### 4.2 Identification of intermediates in the RS-PAH to A-PAH transition

The current work expands our understanding of PAH allostery to include undocked intermediates, simplistically described in Scheme 1.

Scheme 1:

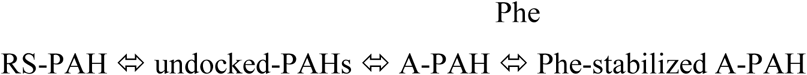

As per the established conformational selection mechanism (15), first suggested by us in 2013 (4), the population of the A-PAH conformation is increased through stabilization by allosteric Phe binding at the ACT dimer interface (circle in Fig 1C). Interestingly, for WT rPAH, the cation-π sandwich at Phe80 appears to act like a lock on the RS-PAH conformation. All tested perturbations lead to an increase in the population of undocked conformers. Possible undocked PAH conformations were recently introduced in a study of the rPAH variant R68S, which is a model for an hPAH disease-associated variant (14). The reported rPAH R68S intrinsic fluorescence, activity, and trypsinolysis results are remarkably similar to what we report herein for the F80L and F80R variants, though the tryptic peptides for R68S were not identified and cleavage within the C-terminal helix was unnoted (14). Included in the characterization of R68S was a small angle X-ray scattering study that was evaluated as an ensemble of undocked structures that allowed flexibility in a linker comprised of residues 110-118. Fitting the data to the undocked conformational ensemble was reported as inferior to an admixture comprised of 33% in the RS-PAH conformation (from crystal structures) and 67% in an A-PAH conformation that held the ACT-domain dimer 8 – 10 Å closer to the tetramer center-of-mass relative to that illustrated in Fig 1C (14). Our ongoing SEC-SAXS studies of rPAH and the Phe80 variants with and without intermediate concentrations of Phe will continue to address undocked models. One such best-fit undocked model is included in Fig 1D (Gupta et al, in preparation); the alternative undocked model in Fig 1D was prepared in conjunction with our ongoing molecular dynamics studies of tetrameric PAH starting in the RS-PAH conformation (Ge, Voelz, et al. in preparation).

### 4.3 Refining A-PAH models

The current work contributes to our building improved models of the A-PAH conformation and provides an improved roadmap towards how to engineer PAH as a structural biology target that is more amenable for study using X-ray crystallography and/or single particle reconstruction using cryo-EM. We initiated this study with a simple model of the PAH quaternary structure equilibrium consisting only of the RS-PAH and A-PAH conformations, positing that destabilizing the former would increase the mole fraction of the latter, thus improving our ability to obtain a crystal structure in the A-PAH conformation. In designing variants to destabilize the RS-PAH conformation, we focused on disrupting the RS-PAH-specific cation-π interaction at Phe80. Unexpectedly our data show that the chosen alterations to Phe80 destabilize both RS-PAH and A-PAH conformations increasing the population of intermediate undocked conformations. Thus, we conclude that Phe80 is important to the stability of both RS-PAH and A-PAH conformations, and posit how Phe80 might stabilize the latter. Our maturing model for the A-PAH conformation considers possible disorder to order transitions. We ask whether disordered segments in the RS-PAH crystal structures become ordered in the A-PAH conformation, and how such disorder ⇔ order transitions might include interactions with Phe80 within A-PAH. Two suggested models are described below.

One model proposes that disordered C-terminal basic sequences in RS-PAH become ordered in A-PAH through cation-π interactions with Phe80. To date, all full-length PAH crystal structures are in the RS-PAH conformation and all are disordered in the C-termini, where the sequence is QKIK(S) (see Fig S1). Each of the proposed A-PAH models (e.g. those shown in Figs 1C and 2B) include juxtaposition of Phe80, on the inner surface of an ACT domain dimer with the C-termini. In the trans conformation shown, the predicted cation-π interaction is inter-subunit; in the cis conformation, it is intra-subunit (13). Arguing against this model is the A-PAH-like character of F80R, where IEC shows that only 10 μM Phe appears to be required to reach the A-PAH conformation. If Phe80 in the A-PAH conformation is interacting with the basic C-terminal residues, F80R should introduce charge repulsion and thus not be A-PAH-like.

A second model considers a disorder to order transition within the N-terminal ∼20 residues (segment 1 of Fig 1A). This segment might move into the space between the ACT domain dimers and the rest of the tetramer. As shown in Fig 1C, this is a fairly large space with an 8-10Å distance between the surfaces of the catalytic domains and the ACT domains. The N-terminal sequence of rPAH is AA**VVLEN**GV**L**S**RKL***S*, where the residues conserved with human PAH are in boldface. The proposed relocation of the N-terminal residues could potentially be stabilized by hydrophobic interactions with Phe80. This second proposed model would have residues ∼1-15 threaded into the space between the ACT domain dimer and the rest of the tetramer with residues ∼18-32 as a potentially disordered linker. In this model residues ∼1-15 go from disorder to order, while residues 18-32 go from order to disorder. We note that Ser16 (italicized above) is a phosphorylation site and phosphorylation is known to reduce the Phe concentration required for full activation (50,51). It is possible that phosphorylation of Ser16 might facilitate the relocation of segment 1 through interaction with Asp112. In support of the notion that the positioning of segment 1 is important to the A-PAH conformation is the reported failure of an rPAH construct missing residues 1-26 to yield an A-PAH crystal structure in the presence of high [Phe] (11).

## 5. CONCLUSION

The current study provides a deeper understanding of the dynamic behavior of mammalian PAH and the profound effect of individual sequence changes at residue 80 on the conformational equilibrium among RS-PAH, A-PAH and undocked intermediate structures. If such undocked structures are relatively more common among disease-associated human PAH variants, this would contribute to the oft-cited generalization that disease-associated proteins are more highly susceptible to degradation (e.g. (33)). Note however that the proteolytic susceptibility of the Phe80 variants is not due to “misfolding”; each of the variant proteins is folded and active. Nevertheless, Phe80 variants are not among the ∼1000 different documented PKU-associated alleles (52). The sum of studies presented herein argues that Phe80 plays a pivotal role in locking PAH into the RS-PAH conformation, potentially providing the rate-determining step in PAH activation. It has long been known that PAH activation is relatively slow and dependent upon the order of addition of reaction components *in vitro* (4,53). Our original interpretation of the unusual kinetic properties of PAH included intermediate dimeric (or monomeric) structures in the pathway between tetrameric RS-PAH and A-PAH conformations, in part to account for the slow timescale of the transition; this remains only a hypothesis (4). The limited proteolysis studies of WT rPAH in the RS-PAH conformation provide strong evidence that tetramer dissociation is facile, consistent with our proposal that reversible multimerization could be a significant part of PAH regulation. As of yet, there has been no systematic inspection of the dynamic oligomerization properties of PKU-associated variants of PAH. Similar to the case of the Phe80 variants reported here, single-residue variants of the wild-type sequence of other proteins, such as a PBGS, have served to successfully predict the behavior and structures of disease-associated and single-residue variants as well as intermediate conformations (26,27,49). We predict that several or many of the ∼1,000 PKU-associated PAH variants affect the dynamic positioning of domains relative to one another as do the Phe80 variants described herein and the PKU-associated R68S.

## Acknowledgements

The authors acknowledge our colleagues Dr. Vincent Voelz of the Temple University Chemistry Department, and Yun-hui Ge for the coordinates illustrated in Fig 1D (right). We also thank Dr. Patrick Loll for his kind gift of the parent plasmid for production of His-SUMO tagged proteins.

## FOOTNOTES

This work was supported by National Institutes of Health Grant 5R01-NS100081 (to E. K. J. and K. G.) and a grant from the National PKU Alliance (to E. K. J.) and in part by the National Institutes of Health NCI Cancer Center Support Grant P30 CA006927. The content is solely the responsibility of the authors and does not necessarily represent the official views of the National Institutes of Health.

The abbreviations used are: IEC, ion exchange chromatography; Phe, phenylalanine; PAH, phenylalanine hydroxylase; rPAH, rat PAH; PKU, phenylketonuria; PAGE, polyacrylamide gel electrophoresis; SDS, sodium dodecyl sulfate; A-PAH, activated PAH; RS-PAH, resting-state PAH; BH_4_, tetrahydrobiopterin; SEC, size exclusion chromatography; SAXS, small angle X-ray scattering; PDB, protein data bank; WT, wild type.

## SUPPORTING INFORMATION FOR

**Figure S1.**
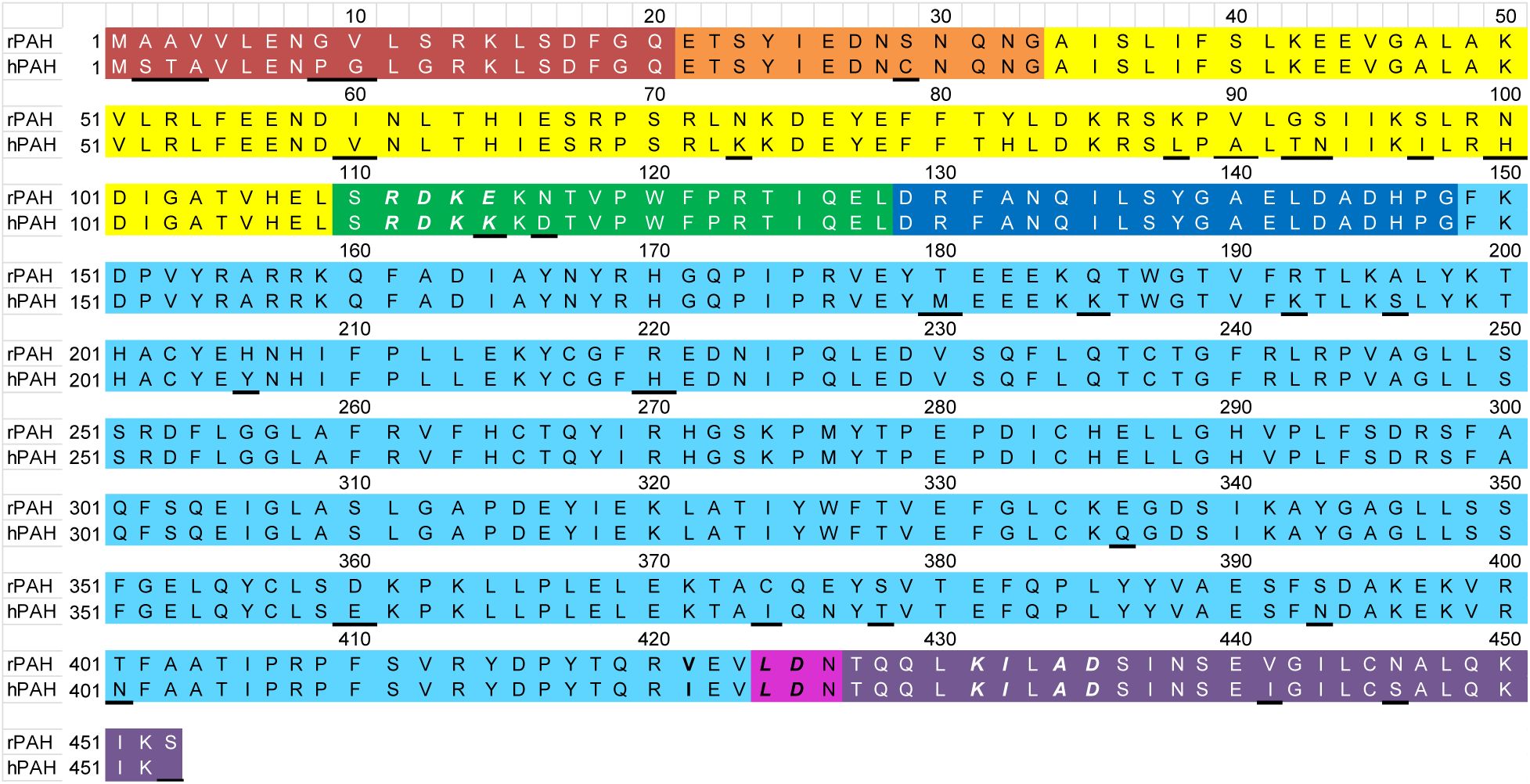
Sequence alignment of WT rPAH and the human (hPAH) proteins. Segments are colored as in Fig 1A. Positions of sequence differences are underlined. Bold italics are used to show the positions of proteolytic cleavage (trypsin and AspN protease) in either the presence or absence of 1 mM Phe.

**Figure S2.**
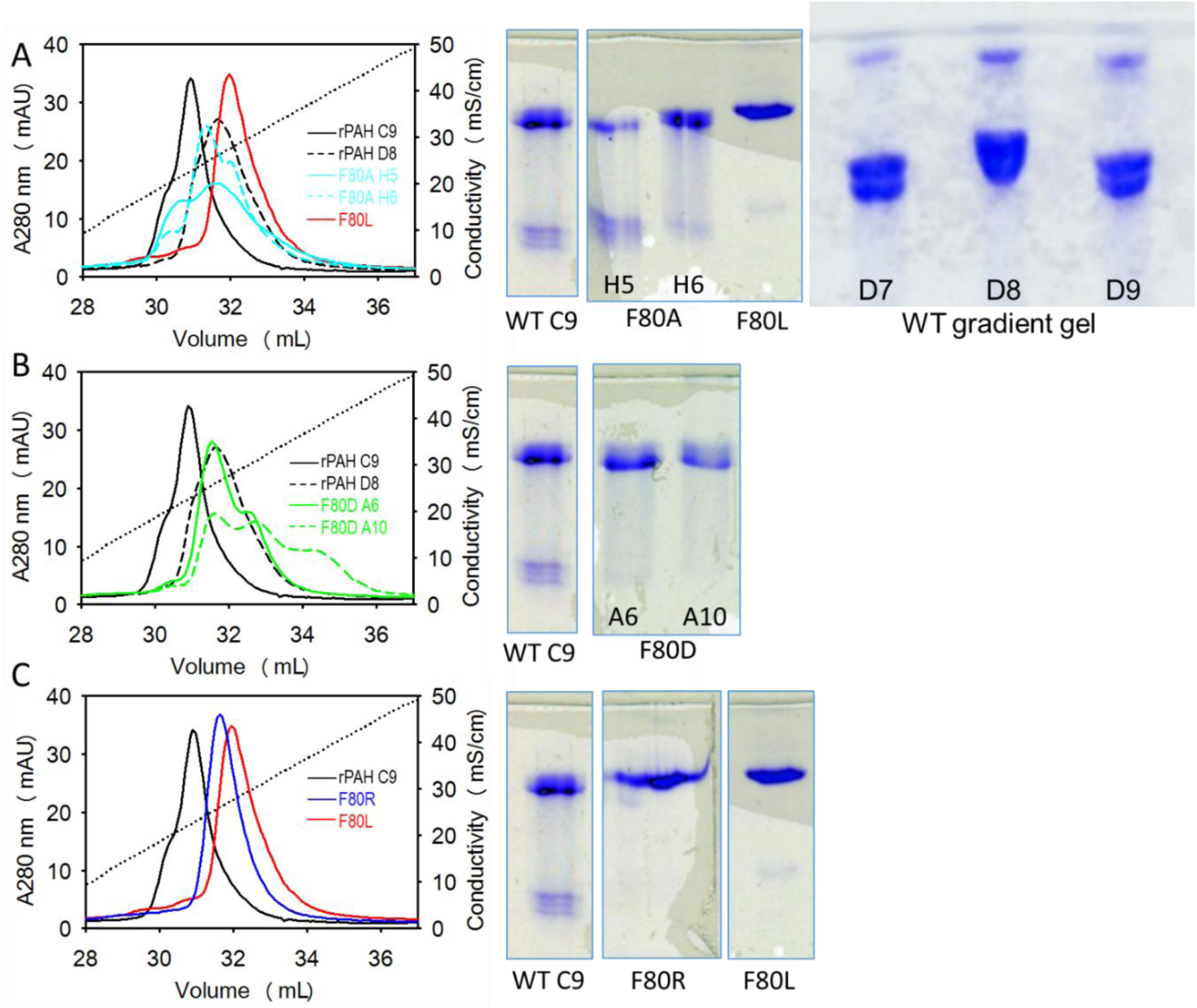
Example analytical IEC and native PAGE of individual fractions from the preparative IEC of WT rPAH and Phe80 variants. For each protein, ∼100 μg of a given fraction was diluted into 5 mL of HiTrap Q buffer at 20 mM KCl (no glycerol), loaded onto and resolved by a 1 mL HiTrap Q column. PhastGels are 12.5%, except where noted. **(A)** Comparison of charge neutral Phe80 variants. The WT rPAH (noted as rPAH) C9 fraction corresponds to the WT rPAH protein for which we determined the full-length RS-PAH crystal structure (1); it is comprised predominantly of the faster migrating tetramer. The WT rPAH fraction D8, from a different preparation, is shown by native PAGE to be comprised of both tetramers. Notably, F80L is resolved from WT rPAH C9 in a way that is similar to what occurs to the retention of WT rPAH upon inclusion of 1 mM Phe (Fig 6c of Ref (2)) (see also Fig 5A). F80A, which is also charge equivalent with WT rPAH, shows multiple tetrameric components, but the earliest to elute does not have the same IEC mobility of either WT rPAH C9 nor of F80L, suggesting that F80A may also be conformationally distinct. **(B)** Comparison of WT rPAH with F80D. **(C)** Comparison of WT rPAH, F80R, and F80L.

**Figure S3:**
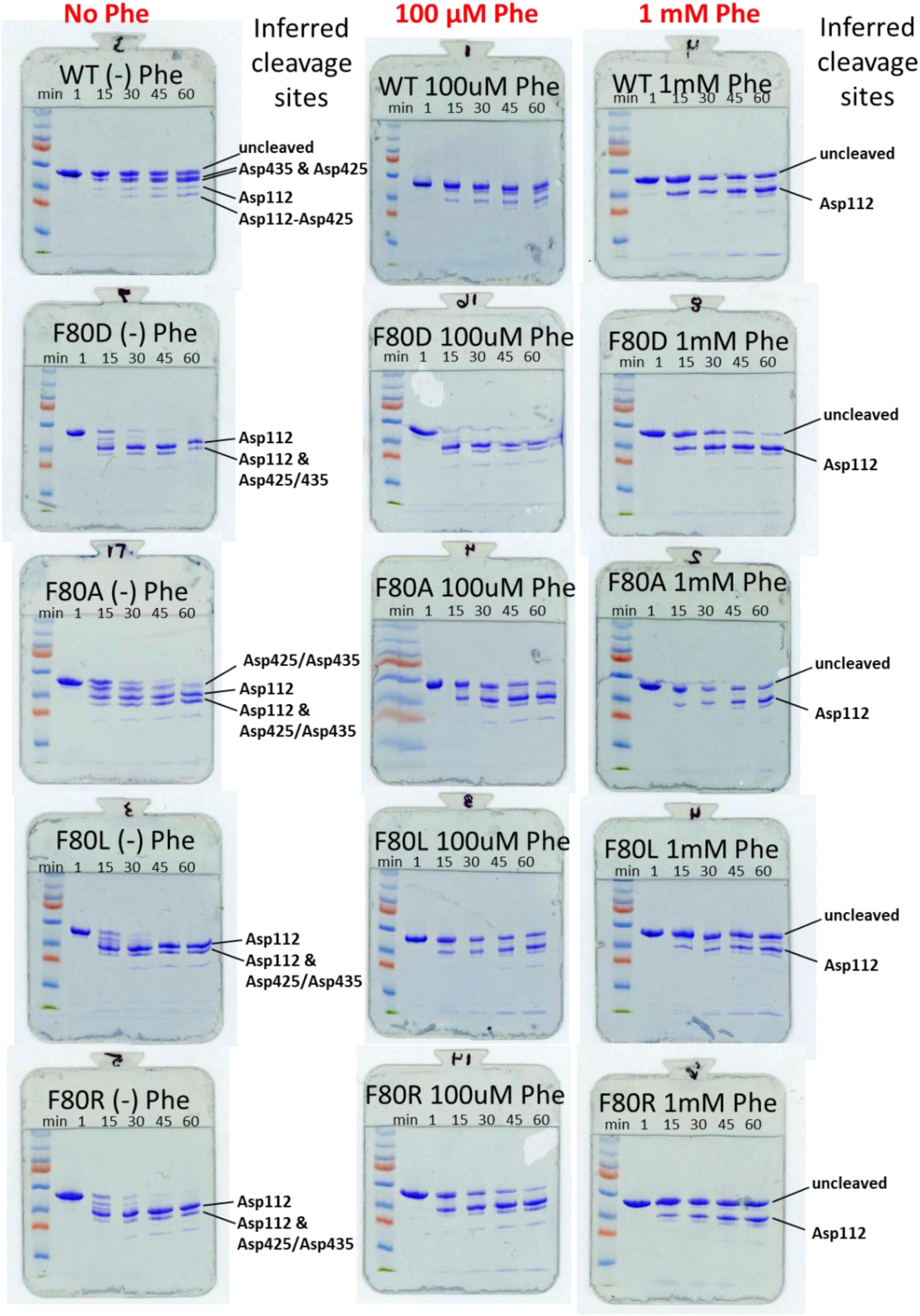
AspN digests of WT rPAH and Phe80 variants as a function of time and of Phe concentration. Each gel is entitled with the protein identity; columns are topped with the concentration of Phe present. The lanes of each gel are labeled with the reaction times of 1min, 15 min, 30 min, 45 min, and 60 min. The leftmost lane is pre-stained molecular weight standards. Identification of cleavage sites is by comparison with Fig 7 of the accompanying paper.

## Notes

### Competing Interest Statement

The authors have declared no competing interest.

### Summary of Updates

The revised manuscript has been submitted for publication in Biochimie, as part of a special issue on inborn errors of metabolism. The manuscript has been substantially reorganized and many of the figures have been revises for improved clarity.

